# Replication of DNA containing trinucleotide repeats by the bacteriophage T7 replisome

**DOI:** 10.1101/2024.11.17.624010

**Authors:** Seung-Joo Lee, Hitoshi Mitsunobu, Sebastian Urbano, Alfredo J. Hernandez

## Abstract

Trinucleotide repeats in the human genome are implicated in various neurodegenerative diseases. The tendency of these repetitive DNA sequences to form non-B DNA structures can cause abnormal replication, leading to genomic instability. This instability contributes to disease progression, though the underlying mechanisms are not fully understood. We investigated the replication of DNA containing CAG and CTG trinucleotide repeats using individual components of the T7 bacteriophage replication machinery, as well as the complete replisome. Our results show that repeats in linear single-stranded DNA (ssDNA) inhibit the activity of T7 DNA polymerase and ssDNA-binding proteins, with a more pronounced effect observed in CTG repeats compared to CAG repeats. Minicircle templates containing CTG repeats exhibited robust DNA synthesis on both the leading and lagging strands, though synthesis was not enhanced by the T7 gene 2.5 ssDNA-binding protein. The lagging strand products generated from the CTG repeat minicircle were significantly longer than those from random sequence templates, and their lengths were not extended by the presence of T7 gene 2.5 protein. When the repeated sequences were incorporated into the T7 phage genome, heterogeneity was observed downstream of the repeats, depending on their length. We propose that aberrant extension occurs predominantly in the lagging strand, driven by dynamic interactions between the repeated sequences and the DNA replisome. This study may provide a foundation for understanding the mechanisms underlying the extension or deletion of repetitive genomic regions.

## Introduction

Many neurodegenerative diseases are associated with unstable genetic elements, such as repeated sequences within gene coding or regulatory regions^1-3^. Although the detrimental effects of these repeated DNA sequences are the result of complex *in vivo* processes involving transcription, DNA repair^4^ and translation^5^, aberrant transmission of genetic information—such as deletion or expansion of repeats during DNA replication— may be a critical initial step in the development of disease^6^. Defective genomic duplication of these repeats can result from the propensity of ssDNA to form non-canonical or alternative DNA structures^7^. During DNA replication, ssDNA generated from unwinding double-stranded DNA (dsDNA) by helicase tends to form stable secondary structures before being used as a template. These non-canonical DNA structures, such as hairpins, cruciforms, and triplexes, can act as roadblocks for DNA polymerase, leading to errors in replication and protein dissociation^8-11^.

Research has focused on understanding the mechanisms by which genomic instability in repeated sequences leads to toxic outcomes. A consensus among findings is that both the location and orientation of these repeats relative to the origin of replication play a critical role in their toxic effects^12^. Another key question is how these repeated sequences are extended during DNA replication. While most studies have focused on cellular-level effects, relatively few have explored the impact of trinucleotide repeats on DNA replication itself. For instance, studies using the T4 phage system have shown that strong secondary structures in the lagging strand can cause separation of DNA polymerase and helicase, resulting in incomplete replication^13-15^. The T7 replisome, with its more comprehensive composition, may provide deeper insights into the replication events involving repeat sequence templates.

The DNA replisome of bacteriophage T7 represents a minimal, yet functional system capable of replicating double-stranded DNA (dsDNA) templates. It consists of T7 gene 4 DNA primase-helicase, T7 gene 5 DNA polymerase, *E. coli* thioredoxin, and the T7 gene 2.5 ssDNA-binding protein, which together coordinate both leading and lagging strand synthesis^16,17^. The interactions between these components and DNA have been extensively studied, making the T7 replisome a valuable model for examining how repeated sequences affect DNA replication^17^.

In this study, we examined the effects of trinucleotide repeats (CTG and CAG) on DNA replication using the bacteriophage T7 replisome. We first assessed the interaction of the repeated templates with individual replication proteins, followed by a comprehensive examination within the full replication complex. We also incorporated these repeats into the T7 genome to investigate how they are maintained in vivo.

### DNA Synthesis Catalyzed by T7 DNA Polymerase on Templates Containing Trinucleotide Repeats

To compare the effects of repeated sequences on DNA synthesis, we constructed linear templates containing 25 repeats of a trinucleotide (CAG or CTG) or random sequences of the same length. These templates included a common sequence where a 15-mer primer annealed to the 3′ region (Fig. 1A). We evaluated the efficiency of nucleotide polymerization by measuring the extension of the ^32^P-labeled primer by T7 DNA polymerase. The amount of full-length 90-base pair (bp) products synthesized on the CAG-repeat template was about 80% of that on the random sequence template, while the CTG-repeat template yielded significantly lower product levels (about 20%) (Fig. 1B).

**Fig. 1.**
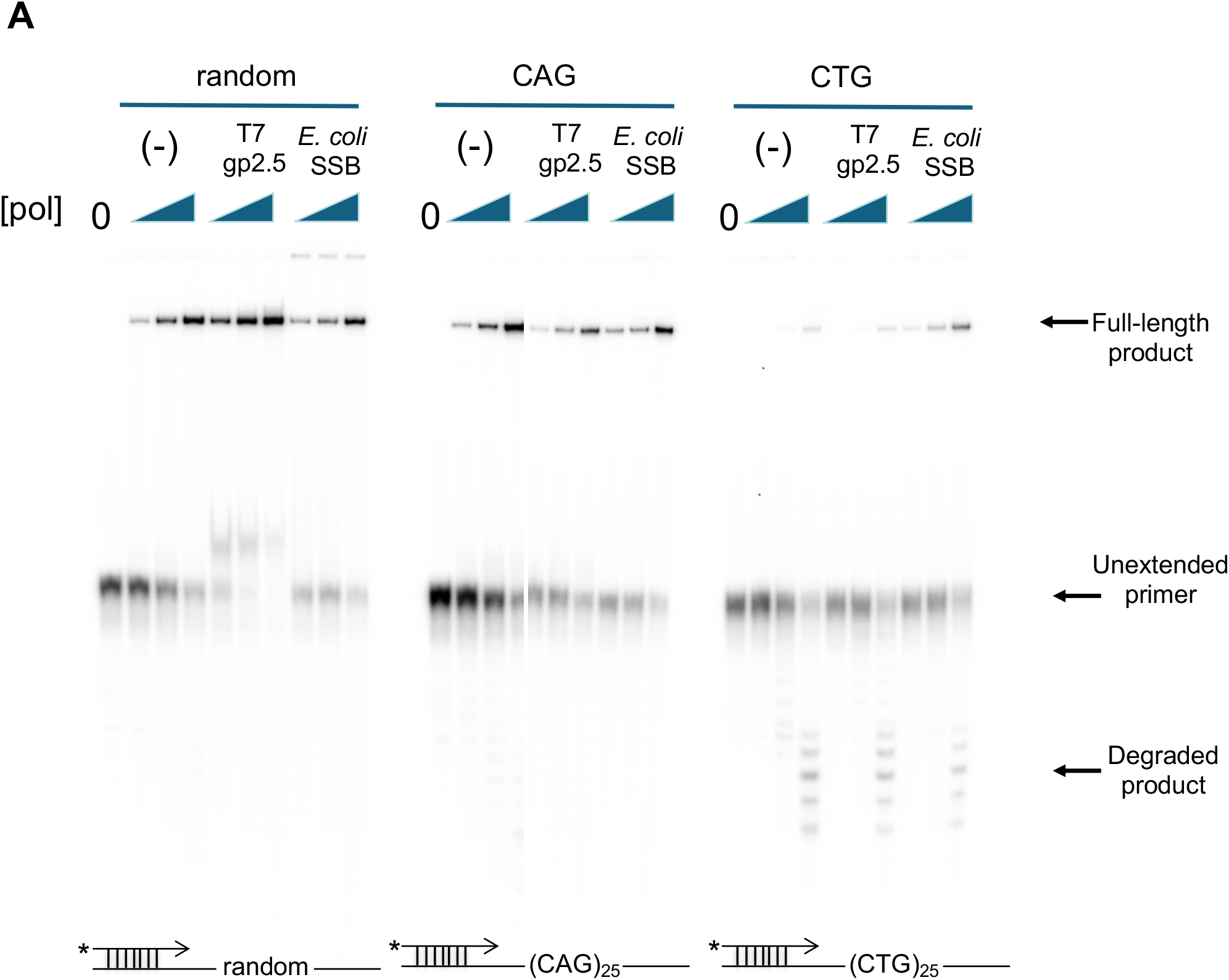
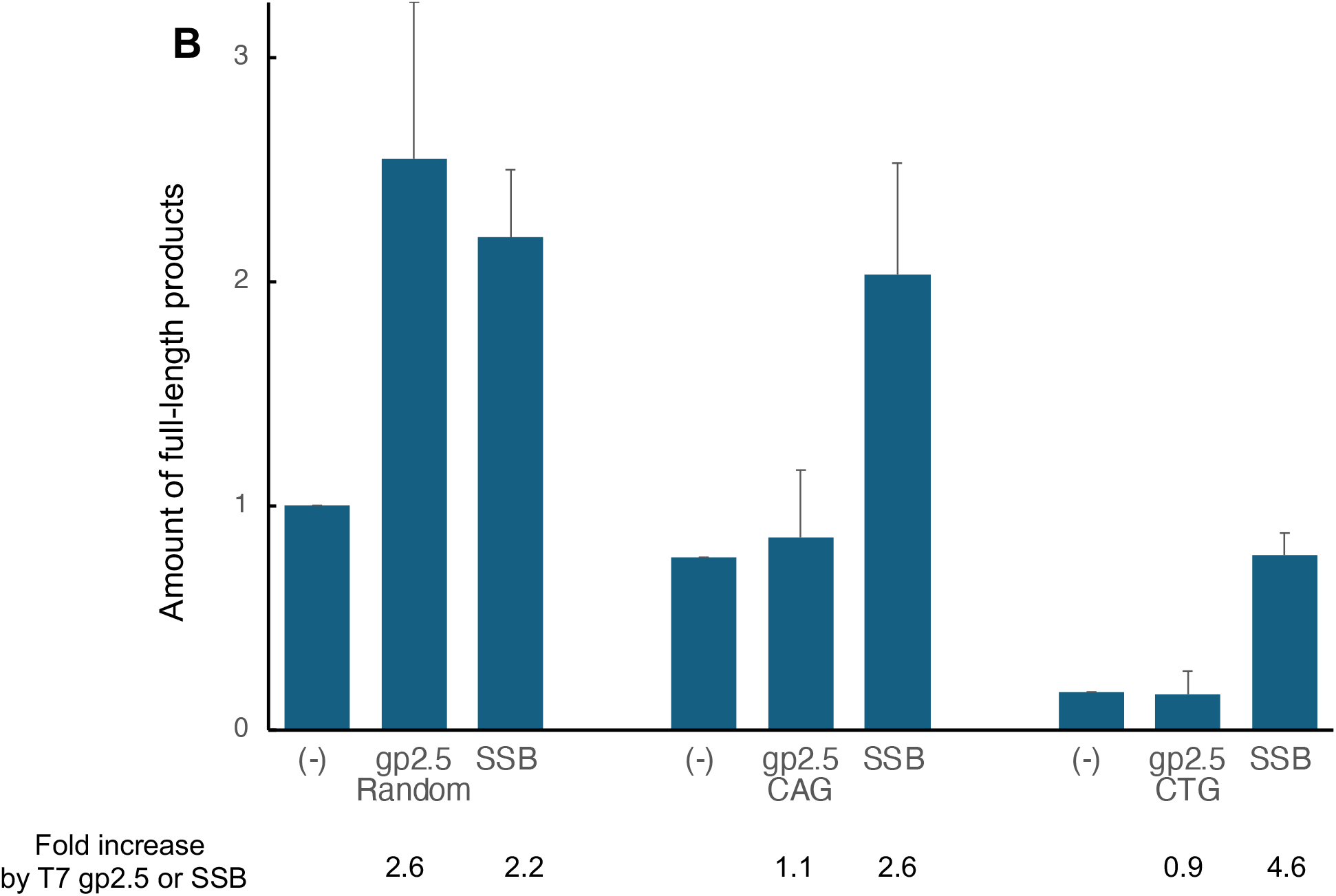
Primer extension on templates containing repeated sequences. (A) A common primer was radiolabeled at the 5′ end and annealed to a template containing either a random sequence or 25 repeats of CAG or CTG sequences. DNA synthesis was performed in standard polymerase reaction mixtures with increasing amounts of T7 DNA polymerase (4, 13, and 40 nM) at 37°C for 3 minutes, in the absence or presence of either T7 gp2.5 or *E. coli* SSB. Reaction products were analyzed on a 20% denaturing polyacrylamide gel. (B) Comparison of full-length products. The amount of full-length products obtained from reactions with 4 nM DNA polymerase was quantified by phosphorimager analysis and normalized against the value obtained with the random sequence template in the absence of additional proteins. The numbers below the columns represent the fold changes resulting from the addition of either T7 gp2.5 or *E. coli* SSB for each set of templates.

Inhibition of DNA synthesis by T7 DNA polymerase was accompanied by excision of newly added nucleotides or degradation of the primer by the enzyme’s proofreading exonuclease activity^18^. Degraded products were observed at the bottom of the gel, particularly with the CTG template, when high concentrations of polymerase were used (Fig. 1A). The reduced levels of full-length products and the presence of more degraded products indicate that DNA synthesis on templates containing repeated sequences is less efficient than on random sequence templates.

Single-stranded DNA-binding proteins can modulate DNA polymerase activity by binding to the DNA template and removing secondary structures that interfere with polymerase progression^19,20^. We assessed the effects of both T7 gene 2.5 ssDNA-binding protein (gp2.5) and *E. coli* ssDNA-binding protein (SSB) on DNA polymerization. Adding T7 gp2.5 increased the yield of fully extended products on random sequence templates by 2.5-fold (Fig. 1B), but no such enhancement was observed with repeated sequence templates. In contrast, *E. coli* SSB enhanced polymerization on all templates by 2 to 5-fold, with the greatest improvement seen on the CTG template (Fig. 1B).

### DNA Binding of T7 gp2.5 and *E. coli* SSB to Linear Templates Containing Trinucleotide Repeats

The inhibition of DNA synthesis by trinucleotide repeats may arise from alternative base pairing or secondary structures that block DNA polymerase progression. The ability of *E. coli* SSB to overcome this inhibition supports this interpretation. However, the differing effects of T7 gp2.5 suggest distinct interactions between the ssDNA-binding proteins and the trinucleotide repeats. To further investigate, we compared the binding affinities of T7 gp2.5 and *E. coli* SSB to 90-mer ssDNA containing random or repeat-containing sequences (Fig. 2). Each 90-mer ssDNA was incubated with increasing concentrations of protein, and the protein-bound DNA was separated from the unbound DNA on a native gel. An 80-mer DNA containing an alternating A/T sequence, known for its high tendency to form secondary structures, was also included in the assay.

**Fig. 2.**
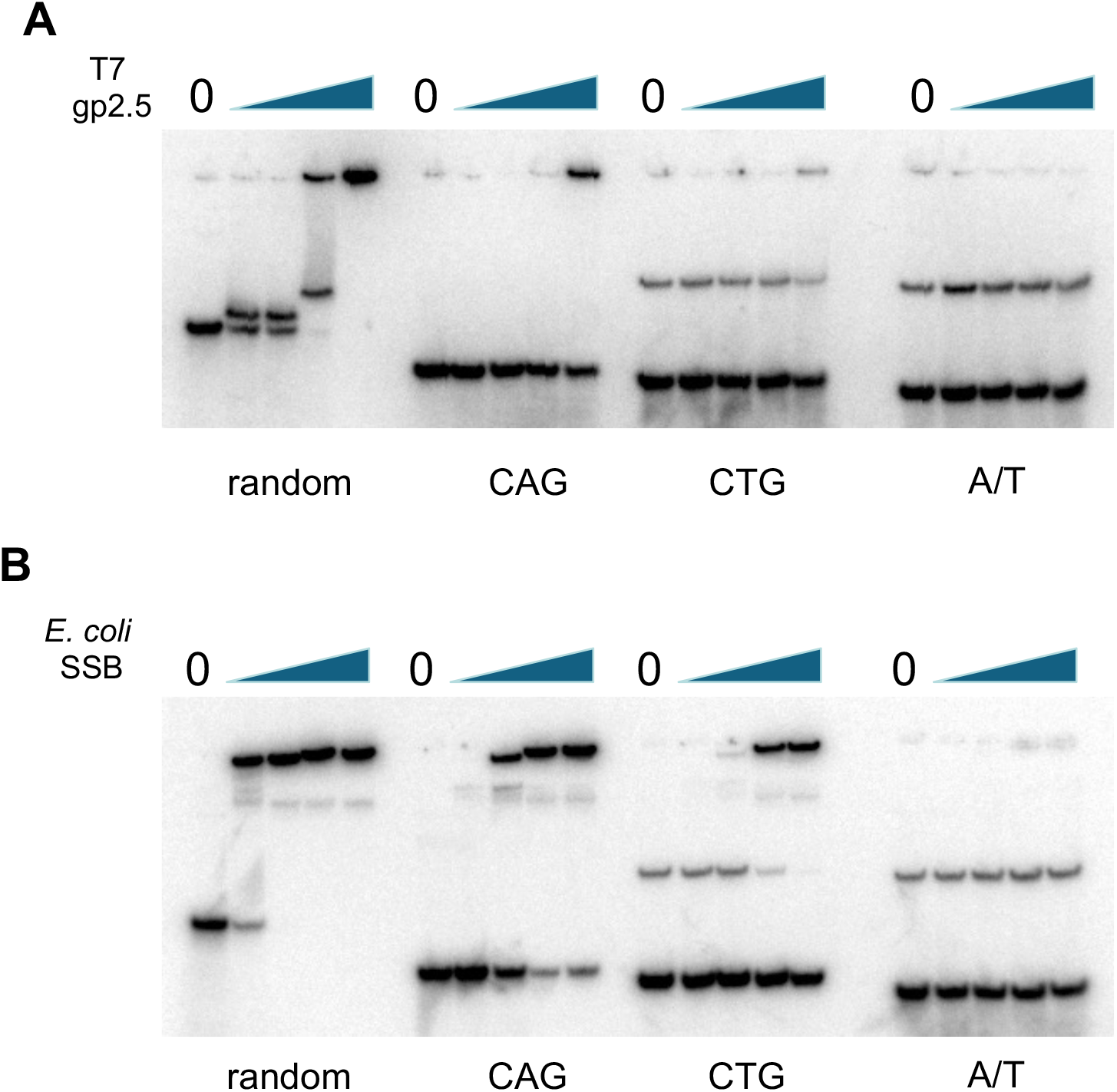
Binding of ssDNA-binding proteins to DNA containing repeated sequences. (A)and (B) show the binding of T7 gp2.5 and *E. coli* SSB, respectively, to ssDNA of various compositions: a 90-mer random sequence, 30 repeats of CTG or CAG, and an 80-mer of alternating A/T sequences. The ssDNA was radiolabeled at the 5′ end and incubated with increasing concentrations (0, 1, 3, 10, and 30 μM) of either (A) T7 gp2.5 or (B) *E. coli* SSB at room temperature for 10 minutes. Protein-bound DNA was separated from unbound DNA using a 10% native polyacrylamide gel electrophoresis conducted at 4°C for 2 hours, followed by visualization through autoradiography.

T7 gp2.5 binds to random-sequence DNA in a concentration-dependent manner but shows minimal binding to DNA containing CAG repeat sequences, except at the highest concentration of gp2.5 (Fig. 2A). DNA containing either CTG repeats or A/T sequences produced a retarded species in the absence of protein, suggesting the formation of secondary structures. However, neither of these sequences exhibited binding with T7 gp2.5. In contrast, *E. coli* SSB binds to DNA containing both CAG and CTG repeats at high protein concentrations, although the binding is significantly weaker than to random-sequence DNA (Fig. 2B). *E. coli* SSB demonstrates a stronger binding affinity to CAG repeat DNA compared to CTG repeat DNA. A similar DNA-binding pattern was observed when primed templates used in the polymerization assays were tested (data not shown).

These results suggest that repetitive sequences are less readily bound by both T7 gp2.5 and *E. coli* SSB, likely due to secondary structure formation. Inhibition of protein binding is more pronounced with CTG repeats than with CAG repeats. Compared to T7 gp2.5, *E. coli* SSB appears to have a greater ability to resolve the secondary structure of repetitive sequences, consistent with its stronger stimulatory effect on DNA polymerization in templates containing repeats (Fig. 1). Both the DNA synthesis and binding assays suggest that the secondary structure of repetitive sequences exerts an inhibitory effect, which is more significant in CTG repeats than in CAG repeats.

### DNA Synthesis on Minicircle Templates Containing Trinucleotide Repeats Design and Preparation of Minicircular Templates Containing Trinucleotide Repeats

A double-stranded minicircle DNA with a 5′ single-stranded tail was employed to monitor both leading and lagging strand DNA synthesis simultaneously^21^. To investigate the effect of repetitive sequences on DNA synthesis, we designed 86-mer minicircles containing 25 repeats of either CAG or CTG, the minimal length reported to be biologically unstable^6^.

Construction of an 86-nucleotide random-sequence minicircle was achieved by ligating the 5′ and 3′ ends using a splint DNA, as previously described^21^. However, this approach did not produce the desired minicircles with repeated sequences, resulting mostly in linear multimers. The secondary structure formation of ssDNA containing repeated sequences likely hindered intramolecular ligation and favored intermolecular ligation between different DNA molecules.

We then used TS2126 RNA ligase I, a thermostable ligase that joins the two ends of the same DNA molecule without a splint DNA^22^, to prepare circular DNA containing either 25 repeats of CAG or CTG. The minicircles were annealed to a 126-mer complementary sequence with a 40-mer poly-T at the 5′ end, resulting in templates containing a common 5′ ssDNA tail and a T7 primer sequence (ACCC) within the circular dsDNA (Fig. 3).

**Fig. 3.**
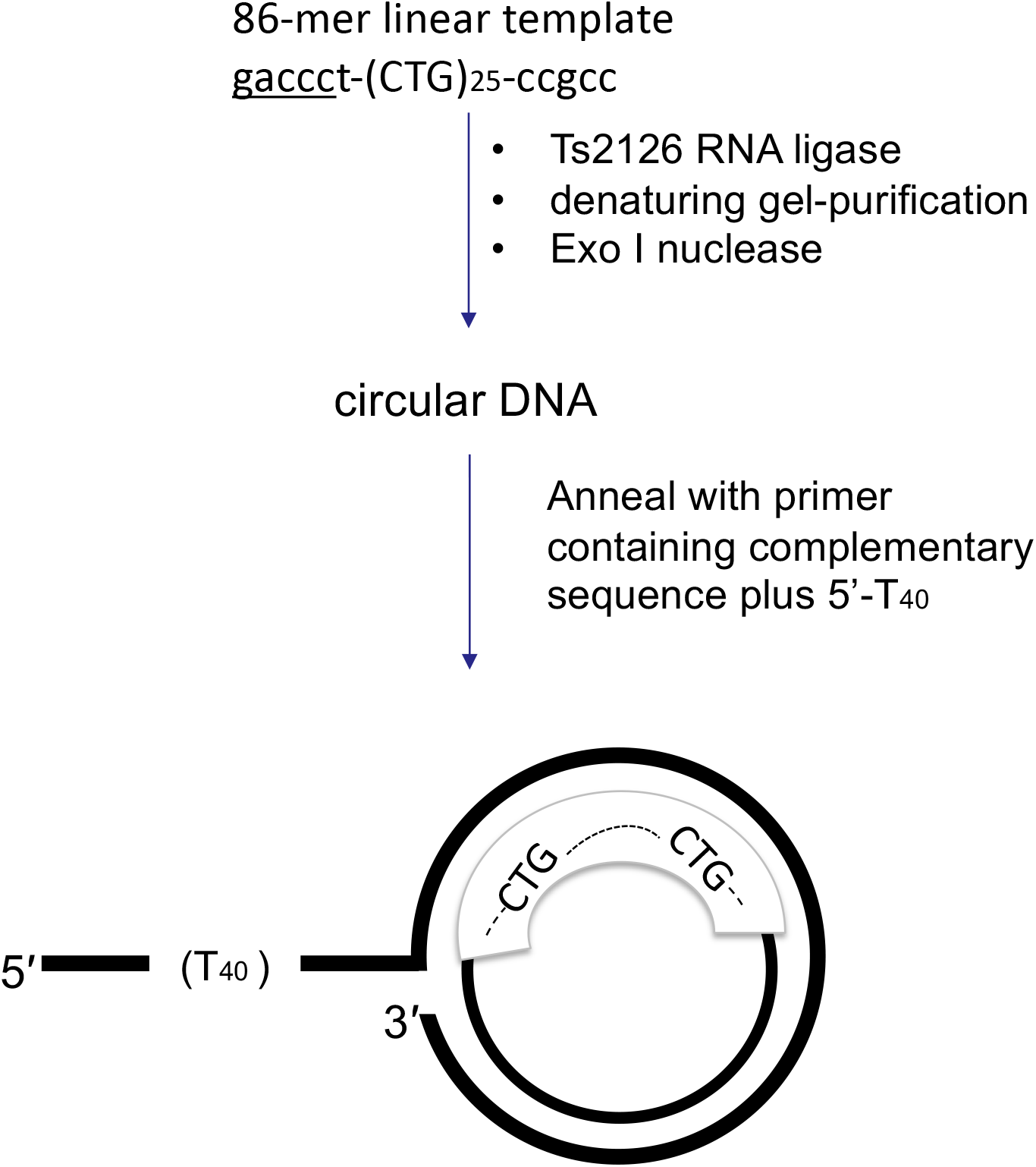
Preparation of minicircular templates containing trinucleotide repeats. A linear 86-mer template DNA containing 25 repeats of either CAG or CTG was circularized using TS2126 RNA ligase. The circular DNA, resulting from intramolecular ligation, was purified using an 8% denaturing polyacrylamide gel and treated with exonuclease I to ensure the removal of any linear DNA. A 126-mer complementary DNA containing a non-complementary T_40_ tail at the 5′ end was annealed to the circular DNA. The schematic representation shows only the CTG minicircular template for simplicity.

Minicircle templates were defined based on the location of the repeat: those with 25 CTG repeats on the circular DNA (the leading strand template) were referred to as CTG repeat minicircle templates, and those with 25 CAG repeats on the leading strand template were referred to as CAG repeat minicircle templates. No significant difference in ssDNA binding by T7 gp2.5 or *E. coli* SSB was observed among the different minicircles.

### Leading Strand DNA Synthesis

DNA synthesis on minicircle templates begins with unwinding by the T7 gene 4 primase/helicase. The helicase domain of the T7 gene 4 protein loads onto the 5′ ssDNA tail, converting dsDNA to an ssDNA template for DNA polymerase, which extends the 3′ end of the primer by incorporating dNMP opposite the circular template (Fig. 3). This continuous strand-displacement process, known as leading strand synthesis, is stimulated by the addition of T7 gp2.5^21,23^.

For minicircles with random sequences, which primarily contain cytosine in the leading strand template, leading strand synthesis was monitored by measuring dGMP incorporation. For CTG and CAG repeat minicircles, the incorporation of dAMP and dTMP, respectively, was measured. Time-course experiments showed that leading strand DNA synthesis on all minicircles was proportional to reaction time (Fig. 4A). The synthesis rate on the random-sequence template was 0.64 µM/min. The CTG repeat template showed a faster rate (1.56 µM/min) compared to the random-sequence template, whereas the CAG repeat template had a similar rate (0.6 µM/min). Adding T7 gp2.5 increased the amount of dNMP incorporated, enhancing the rate by approximately 3-fold in both random and CAG templates, but no similar enhancement was observed with the CTG template (Fig. 4A, Table 1).

**Table 1.**
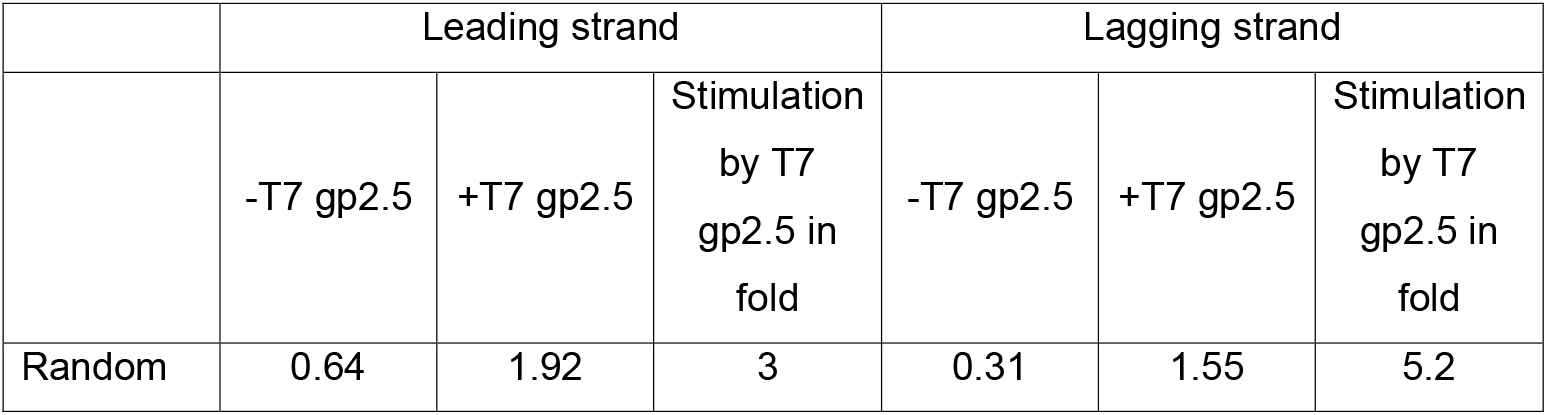
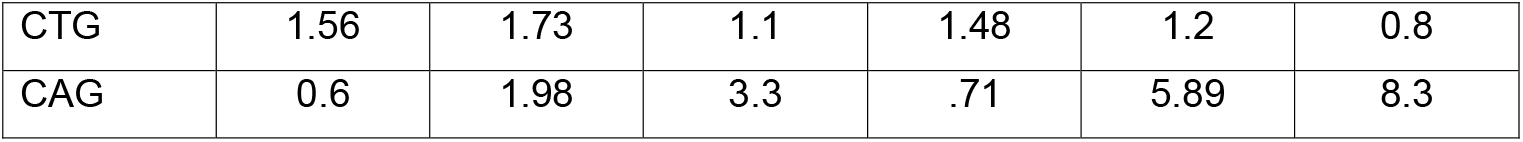
Rates of DNA synthesis on minicircle templates Slopes of plots shown in Fig. 4 was calculated and presented in µM dNMP incorporated/min. Stimulation by T7 gp2.5 was obtained by dividing the synthesis rate in the presence of gp2.5 by the synthesis rate in the absence of gp2.5.

|  | Leading strand |  |  | Lagging strand |  |  |
| --- | --- | --- | --- | --- | --- | --- |
|  | -T7 gp2.5 | +T7 gp2.5 | Stimulation<br>by T7<br>gp2.5 in<br>fold | -T7 gp2.5 | +T7 gp2.5 | Stimulation<br>by T7<br>gp2.5 in<br>fold |
| Random | 0.64 | 1.92 | 3 | 0.31 | 1.55 | 5.2 |
| CTG | 1.56 | 1.73 | 1.1 | 1.48 | 1.2 | 0.8 |
| CAG | 0.6 | 1.98 | 3.3 | .71 | 5.89 | 8.3 |

**Fig. 4.**
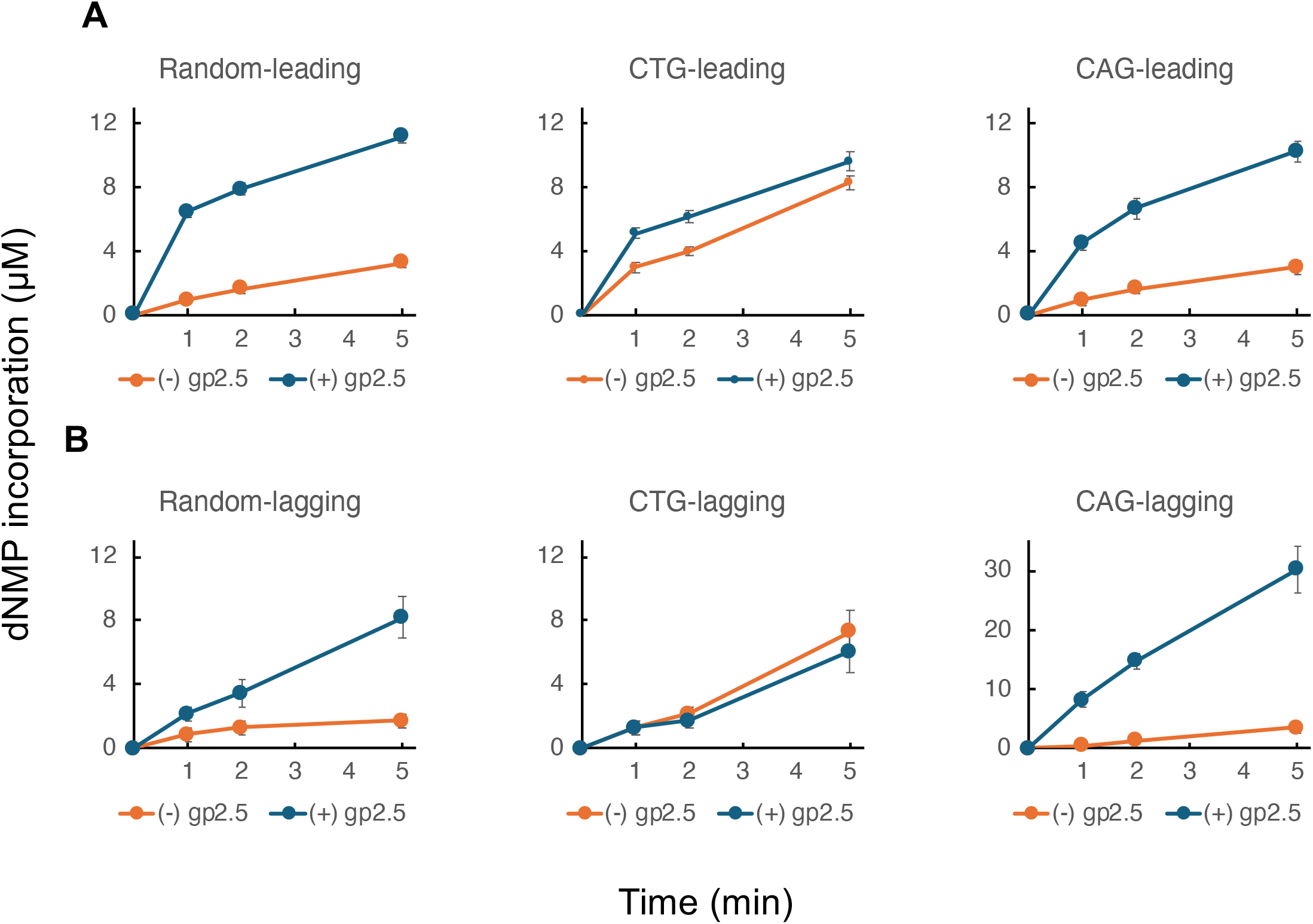
Leading and lagging strand DNA synthesis on minicircles containing random and repeated sequences. DNA synthesis was determined by measuring the incorporation of specific radioactive nucleotides (indicated in parentheses for each strand) present almost exclusively in each strand of the minicircle templates. The reaction mixtures contained 300 μM each of ATP and CTP, 500 μM of each dNTP including the [α-^32^P] dNTP specific for leading or lagging strand synthesis, 10 nM minicircle template, 80 nM T7 DNA polymerase, 60 nM T7 gene 4 primase-helicase, and 4 μM T7 gene 2.5 protein. Reactions were performed at 30°C for the indicated times. To normalize DNA synthesis rates across templates of different lengths, all rates of DNA strand synthesis on the repeated sequence templates were adjusted by multiplying by a factor of 45/26 = 1.7, since the random sequence template contained more measurable nucleotides (45 dGMP for leading strand and 45 dCMP for lagging strand) compared to the repeated templates (26 dAMP or dTMP).

### Lagging Strand DNA Synthesis

As leading strand synthesis progresses, ssDNA is extruded behind the helicase (Fig. 3). The primase domain of T7 gp4 generates tetraribonucleotides at primase recognition sites on the ssDNA^24^, serving as primers for a second DNA polymerase to initiate lagging strand synthesis. The addition of T7 gp2.5 stimulates lagging strand synthesis. Lagging strand synthesis on the random-sequence template was slower (0.31 µM/min in the absence of gp2.5, 1.55 µM/min in the presence of gp2.5) than leading strand synthesis (Fig. 4B, Table 1). The rate of lagging strand synthesis on the CTG repeat template without gp2.5 was 5-fold higher than that of the random-sequence template.

The rate on the CAG repeat template was also higher than on the random-sequence template but lower than on the CTG repeat template (Table 1). The addition of T7 gp2.5 increased the lagging strand synthesis rate in both the random (3-fold) and CAG repeat templates (8-fold), but no stimulation was observed with the CTG template.

### Effect of T7 gp2.5 on Leading and Lagging Strand DNA Synthesis

T7 gp2.5 is crucial for coordinated DNA synthesis, where leading and lagging strand syntheses are enhanced and occur at similar rates^21,23^. In a previous study using a 70-mer minicircle, a range of T7 gp2.5 concentrations (2-6 µM) supported coordinated synthesis, with no effect beyond 2 µM^21,23^. In our study, 4 µM T7 gp2.5 stimulated DNA synthesis in most templates except the CTG template (Fig. 4), but the rates of leading and lagging strand synthesis did not appear coordinated when using CTG and CAG containing minicircles as templates. To explore this further, we tested a range of T7 gp2.5 concentrations on various templates (Fig. 5).

**Fig. 5.**
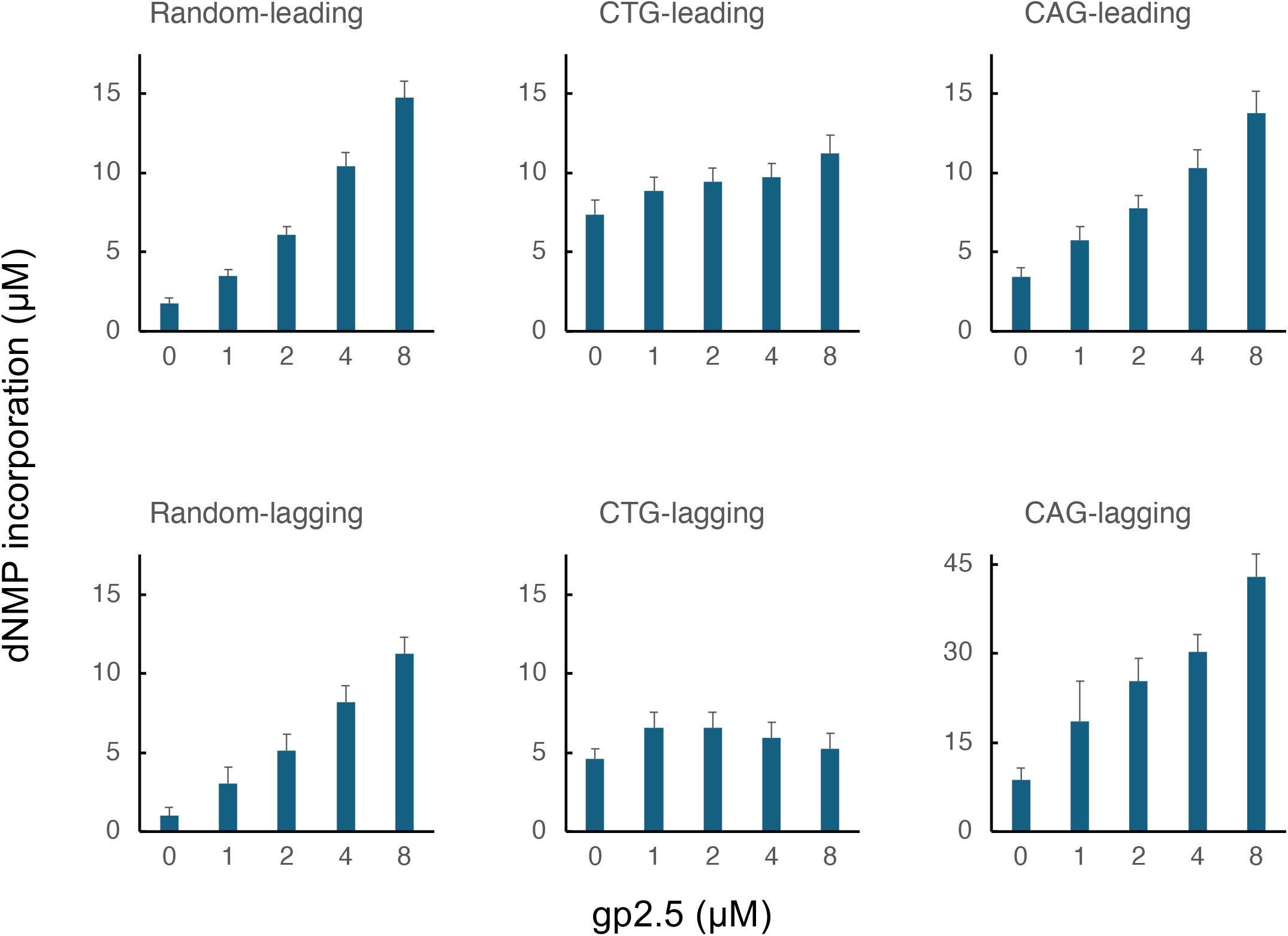
Effect of T7 gp2.5 concentration on leading and lagging strand DNA synthesis of minicircles containing random or repeated sequences. DNA synthesis was measured by the incorporation of dNMP into DNA for each template, as described in Fig. 4. Reactions were carried out at 30°C for 5 minutes with T7 gp2.5 concentrations of 0, 1, 2, 4, and 8 μM.

For the random-sequence template, both leading and lagging strand syntheses were stimulated proportionally to the amount of T7 gp2.5, showing an almost 10-fold increase at the highest concentration. While leading strand synthesis on the random template increased over 10-fold at the highest concentration, it was less than 2-fold for the CTG template. Stimulation of leading strand synthesis on the CAG template was less pronounced than on the random template but increased up to 4-fold at the highest T7 gp2.5 concentration. Lagging strand synthesis on the CTG template did not correlate with the amount of T7 gp2.5 added. The addition of gp2.5 stimulated lagging strand synthesis on the CAG template by 5-fold, but the amount of DNA synthesized was much higher than on the random-sequence template.

### Analysis of DNA Products on Alkaline Agarose Gel

DNA products from the leading and lagging strand synthesis assays were analyzed by alkaline agarose gel electrophoresis (Fig. 6). The leading strand products for all templates in the absence of gp2.5 were relatively long, ranging from 2 to 9 kb, with CAG template products slightly longer than those from other templates. Stimulation by gp2.5 was most evident for the random-sequence template, consistent with previous results (Figs. 4 and 5).

**Fig. 6.**
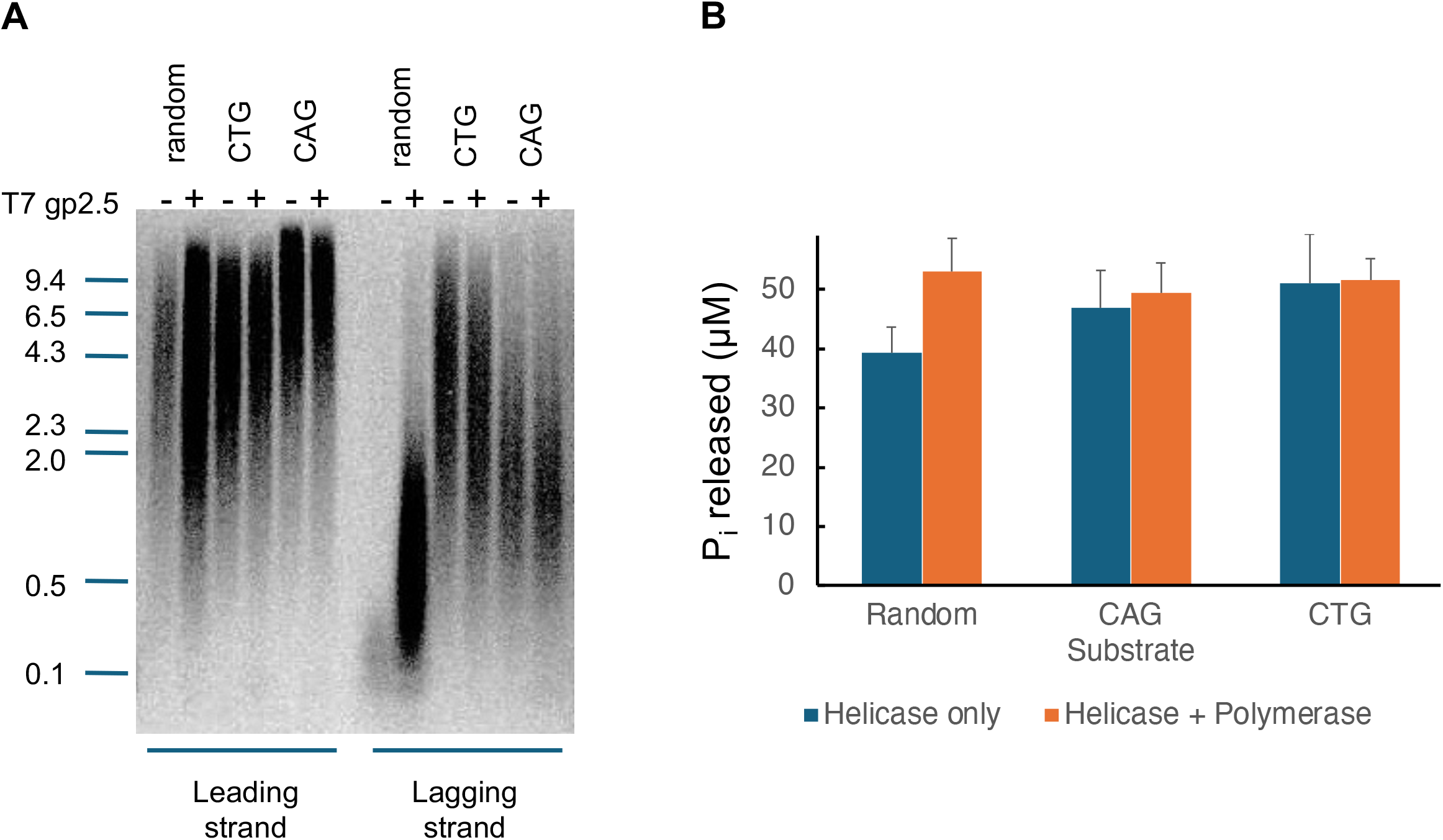
(A) Analysis of DNA synthesis reaction products from minicircle templates containing random and trinucleotide repeat sequences. Reactions were performed for 5 minutes under the same conditions as described in Fig. 4. After the removal of unincorporated radioactive dNTPs, reaction products were loaded onto a 0.8% alkaline agarose gel and run overnight. The gel was dried and visualized by autoradiography. (B) dNTP hydrolysis in the presence of minicircles containing random and repeated sequences. Reactions were incubated for 5 minutes and released phosphate was quantified by the colorimetric malachite green assay.

Lagging strand synthesis showed a more pronounced difference in product length. Consistent with prior studies^21,23^, lagging strand products from the random-sequence template were short (∼0.2 kb) without gp2.5 and increased up to 2 kb with gp2.5 addition. In contrast, lagging strand products from trinucleotide repeat templates were much longer in the absence of gp2.5, and their lengths were unaffected by gp2.5. Products from the CTG repeat template were notably longer than those from the CAG repeat template.

### Replication of Trinucleotide Repeats in the T7 Genome

The *in vitro* results demonstrate that trinucleotide repeats have a significant impact on DNA synthesis within the reconstituted T7 replisome. To investigate the effect of these sequences on DNA replication *in vivo*, we introduced random, or trinucleotide repeat sequences into the T7 phage genome using homologous recombination.

A 1 kb DNA fragment containing the sequences of interest was inserted into the pUC19 vector at a *Sma I* site. This fragment spanned a region of the T7 genome surrounding gene 5.3, a non-essential gene, that was replaced with the *E. coli* thioredoxin gene, immediately followed by an 86-mer containing either a random sequence or trinucleotide repeats (Fig. 7A). *E. coli* cells containing the plasmid were infected with wild-type T7 phage, and recombinant phage from the cell lysate was collected. Since *E. coli* thioredoxin is essential for T7 DNA replication and phage growth, phages that successfully acquired the inserted DNA (containing the thioredoxin gene) were selected by growing them on *E. coli* lacking thioredoxin.

**Fig. 7.**
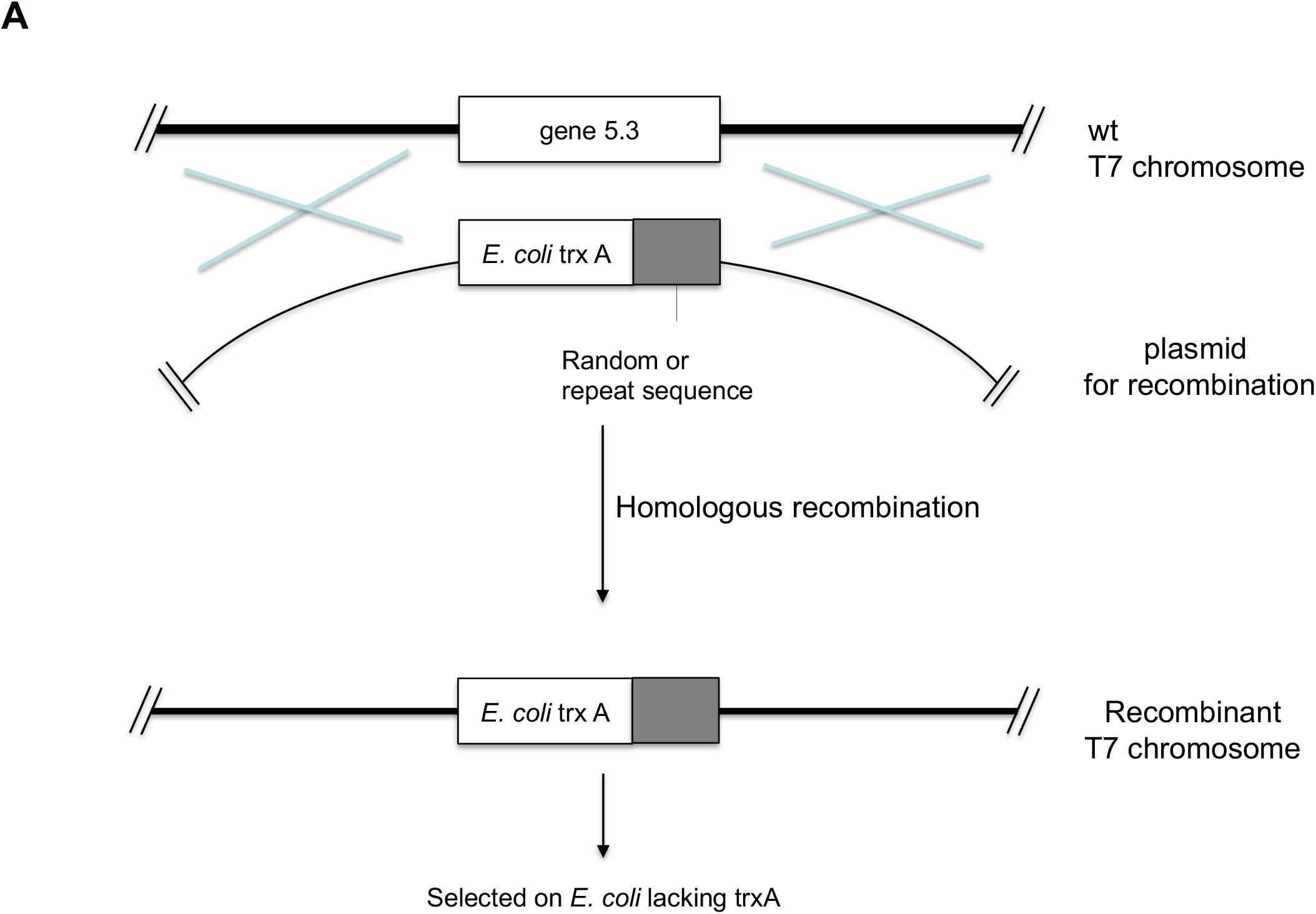
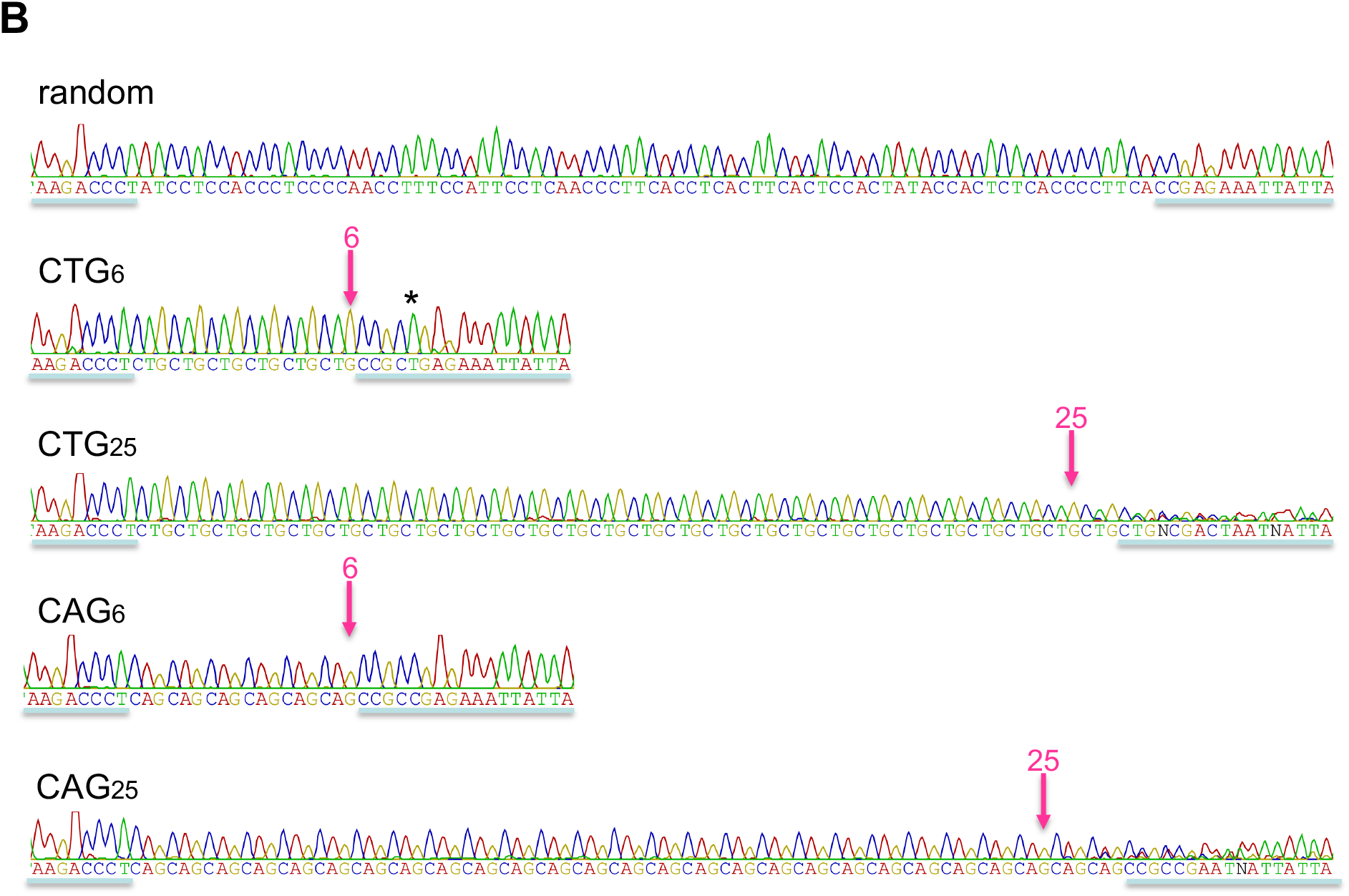
Replication of trinucleotide repeats in the T7 phage genome. (A) Introduction of random and trinucleotide repeat sequences into the gene 5.3 locus of the T7 phage genome by homologous recombination. The plasmid used for recombination contains the *E. coli* thioredoxin gene, immediately followed by an 86-mer of either random or trinucleotide repeat sequences. Phages that successfully replaced gene 5.3 with the inserted DNA were selected by growing on *E. coli* lacking the thioredoxin gene. (B)DNA sequence analysis of inserted trinucleotide repeats in the T7 genome. The recombined region was amplified by PCR, and the DNA sequence was determined for the inserted random sequence, 6 repeats of CTG, 25 repeats of CTG, 6 repeats of CAG, and 25 repeats of CAG. The common regions flanking the random or repeat sequences are underlined. The downstream end of the repeated sequences from the initial plasmid is marked with a red arrow. A single substitution (C to T, denoted by *) in the flanking region of the CTG_LJ_ repeat originated from the initial plasmid before recombination.

The inserted sequences included the same random sequence, a 25-repeat CAG, or a 25-repeat CTG as used in the minicircle studies. Additional inserts with shorter repeats, obtained during plasmid construction (6, 8, and 12 repeats of CTG; 6, 9, and 17 repeats of CAG), were also incorporated into the T7 genome. Recombinant phages were isolated from single plaques and amplified by infecting *E. coli* lacking the thioredoxin gene. The genomes of the recombinant phages containing random or repeated sequences were then amplified by polymerase chain reaction (PCR) using diluted phage lysates as templates, followed by DNA sequence analysis.

The recombination frequency, as indicated by the number of plaques resulting from successful recombination, was similar across all tested sequences. DNA sequence analysis of recombinant phages showed that the random sequence was faithfully amplified in the phage genome without modification (Fig. 7B). Short repeats (6 repeats of both CTG and CAG) were also accurately amplified. However, longer repeats (25 repeats of both CTG and CAG) resulted in ambiguous sequences downstream of the repeat region, suggesting the presence of mixed phage populations. Although some sequences shown in Fig. 7B appeared to extend beyond the original 25 repeats, shorter repeats were more frequently observed in both CAG and CTG repeat phages. Similarly, recombinant phages containing a relatively long repeat (17 repeats of CAG) also exhibited sequence heterogeneity (Table 2), indicating that longer repeats are prone to producing heterogeneous progeny.

**Table 2.** Heterogeneity of repeated sequence in recombinant phages DNA from recombinant phages containing different lengths of either CAG or CTG repeat was examined for sequence analysis. – indicates that DNA analysis shows homogeneous DNA sequence. + denote that DNA analysis demonstrates ambiguous DNA sequence downstream of repeats presumably due to existence of heterogeneous species. At least two independent phage plaques were tested for each length of repeat.

|  | (CTG)<sub>n</sub> | (CAG)<sub>n</sub> |
| --- | --- | --- |
| n= 6 | – | – |
| 8 | – |  |
| 9 |  | – |
| 12 | – |  |
| 17 |  | + |
| 25 | + | + |

The observed variation in DNA sequences likely arises during the amplification of the phage genome after recombination since homologous recombination occurs through the flanking sequences of the repeat and all tested sequences should integrate into the T7 genome with similar efficiency.

## Discussion

We examined DNA replication of repeated sequences by first investigating their interactions with individual T7 DNA replication proteins and then with the complete T7 replisome using a minicircle template. We found that secondary structure formation in linear single-stranded DNA (ssDNA) with repeated sequences impedes primer extension by T7 DNA polymerase and binding by the T7 gene 2.5 ssDNA-binding protein (gp2.5). This inhibitory effect was more pronounced with CTG repeats than with CAG repeats. While *E. coli* SSB protein binds and enhances primer extension by T7 DNA polymerase regardless of the template sequence, T7 gp2.5 stimulated polymerase activity only on the random sequence template. Our results using a minicircular DNA template with the full T7 phage replisome revealed several key observations: **1)** DNA synthesis on the CTG template, in the absence of T7 gp2.5, was much higher than on the random sequence template for both the leading and lagging strands. **2)** The level of DNA synthesis stimulation by T7 gp2.5 was similar for both the random and CAG templates, but no stimulation was observed for the CTG template. **3)** The DNA synthesis products from repeated sequence templates were significantly larger than those from the random sequence template, especially on the lagging strand. While the random template initially produced short lagging strand segments, which were extended significantly with the addition of T7 gp2.5, no such changes were observed with either repeated sequence template.

The tendency of repeated sequences to form secondary structures in linear DNA correlates with their inhibitory effects on most replication proteins, with a stronger effect seen in CTG repeats than in CAG repeats. This finding aligns with previous observations that more stable secondary structures provide greater inhibitory effects on replication^25^. However, the results from the minicircle template are more complex than what could be explained by a simple inhibitory effect. Not only do replication proteins interact differently with repeated sequences, but the various protein components of the replisome also interact with each other, influencing their activities. For example, it has been shown that the speed of gene 4 helicase is enhanced by its interaction with DNA polymerase at the replication fork during strand-displacement DNA synthesis^26^. Consistent with our observations using individual replication components, the gene 4 helicase alone translocates or unwinds DNA containing repeated sequences at a slightly higher rate compared to random sequences, as measured by dNTP hydrolysis (Fig. 6B). However, when coupled with T7 DNA polymerase for strand displacement synthesis, no significant difference in dNTP hydrolysis was detected among the different template sequences. This suggests that the helicase, when complexed with DNA polymerase, unwinds the minicircle template equally regardless of sequence context.

A striking observation with the repeated templates was the significant enhancement of DNA synthesis on the CTG template. How is this increased synthesis achieved? As mentioned earlier, the rate of DNA unwinding by the T7 gene 4 complexed with T7 DNA polymerase appears to be equivalent for all templates. Thus, increased incorporation of dNMP on the CTG template could be due to either preferential incorporation by DNA polymerase or the generation of ssDNA templates that allow for preferential primer extension. Given that the rate of dNMP incorporation by T7 DNA polymerase does not vary significantly for all nucleotides^18,27^, the former possibility is unlikely. Instead, the unwound CTG template region must remain at least temporarily free from secondary structure, allowing DNA polymerase to incorporate cognate nucleotides and extend the nascent CAG strand. If the exposed ssDNA CTG template at the fork is long enough to temporarily stabilize itself by forming a secondary structure after polymerization, it will prevent the growing CAG strand from base pairing. In this scenario, the tendency of the ssDNA CTG template to form stable structures could drive faster synthesis of the leading strand. Alternatively, the conformation generated in the template strand might facilitate DNA polymerization by stabilizing the binding of DNA polymerase. Enhanced levels of leading strand synthesis were observed only with the CTG template, which has a high intrinsic propensity to form secondary structures. A detailed understanding of the active replication fork structure would be necessary to confirm this hypothesis.

For the lagging strand, the template extruded from the fork easily forms secondary structures until T7 DNA primase initiates de novo synthesis of oligonucleotides. Although a primase recognition site is present in the common sequence of all templates, only the one in the repeated templates likely remains in ssDNA form due to secondary structure formation, providing better access for T7 primase to initiate lagging strand synthesis. The lengthy lagging strand synthesis products from the repeated templates suggest that the source of enhanced synthesis might be longer extension products rather than frequent priming. Similar to what happens on the leading strand, the tension created by secondary structure formation within the replication bubble might facilitate rapid polymerization by stabilizing the ssDNA region near the polymerization site. This rapid movement of DNA polymerase could cause the primase to skip recognition sites in the template, resulting in longer products. It has been previously suggested that polymerase skipping over folded-back templates in repeated sequences could lead to repeat deletions. This fold-back deletion or shortening of repeats could be explained by secondary structure formation in the template before polymerization occurs, as described in the slippage model.

T7 gp2.5 has been shown to enhance and coordinate leading and lagging strand synthesis in the minicircle template^21^. The minicircle template containing the CAG repeat showed a similar level of increased synthesis to the random sequence template by T7 gp2.5 (Table 1). In contrast, the minicircle template containing the CTG repeat did not show significant enhancement by T7 gp2.5, particularly for the leading strand. This suggests that the effect of secondary structure on leading strand extension is critical for its interaction with T7 gp2.5. Although the CTG repeat, generated the nascent leading strand products of the CAG repeat minicircle, is also prone to forming secondary structures, it behaves differently from the leading strand synthesis on the CTG repeat minicircle. Notably, T7 gp2.5 shows weak or no binding to ssDNA of repeated sequences, particularly the CTG repeat (Fig. 2). Presumably, in the context of the replisome, T7 gp2.5 binding to repeats may be mediated by other factors, resulting in different affinities. Unlike previous studies using a 70-mer minicircle template^21,23^, the rate of DNA synthesis for leading and lagging strands was not equal in CTG and CAG-containing minicircles. Nevertheless, the stimulation of DNA synthesis by T7 gp2.5 and the interdependence of leading and lagging strand syntheses suggest that they are at least partially coordinated under our conditions.

The minicircle DNA used in this study offers several advantages over other templates. First, it allows simultaneous monitoring of both leading and lagging strand DNA synthesis. Since the linear template for the lagging strand is produced by strand displacement DNA synthesis during leading strand synthesis, measuring lagging strand synthesis also indicates that leading strand synthesis has occurred beforehand or simultaneously, enabling the monitoring of both strands. Second, we have the flexibility to design the DNA sequence of the minicircle template, allowing us to enrich the content of the sequences to be examined. In the trinucleotide repeat template, we included 25 repeats (25 nt x 3 = 75 nt) and an 11 nt non-repeated common sequence containing a T7 primase site, making almost 90% of the minicircle sequence consist of the repeated sequence. This high content of repeated sequences makes it easier to determine their effects on DNA synthesis. The difficulties in preparing minicircle templates containing repeated sequences due to secondary structure formation were overcome by using TS RNA ligase. These merits of the minicircle template enable us to effectively analyze replication on repeated templates.

The *in vivo* assays, in which repeated DNA sequences were integrated into the T7 genome, indicate that longer repeated sequences are not faithfully maintained. We expected that the insertion of the CTG repeat would cause more aberrant replication than the others, given that both the leading and lagging strands of the CTG minicircle template showed drastic differences from the other templates. However, we observed that both CAG and CTG repeats produced heterogeneous sequences downstream of the inserted sequence, suggesting that the longer extended DNA lagging strands generated from both repeated templates might be responsible for the irregular replication. Despite the lack of sequence preference observed *in vivo*, we cannot rule out that the severity of CTG replication was underestimated due to the limited sensitivity of phage genome analysis. For sequence analysis, we used phage lysate that had already undergone multiple rounds of propagation after the introduction of repeats into the T7 genome. The CTG repeat could have been more prone to sequence length abnormalities earlier than the CAG repeat, but this difference might have diminished during multiple amplification steps. It has been reported that a minimum of 25 trinucleotide repeats is necessary to observe genome instability^28^. Unlike other systems, a shorter repeat length (as short as 17) was sufficient to show genome instability in the streamlined T7 phage replication system.

The propensity of trinucleotide repeats to form intermolecular base-pairing affects their interactions with the protein components of the replisome. The enhancement of DNA synthesis on the CTG repeat template in the absence of T7 gp2.5, along with the lengthy extension of lagging strand synthesis, highlights the abnormalities of DNA synthesis from repeated templates. The dynamic nature of macromolecular interactions between replisome components and repetitive sequences is likely a major contributing factor for pathological replication of repeated sequences and contributes to genomic instability.

### Experimental Procedures

#### Materials

All synthetic DNAs were obtained from Integrated DNA Technologies. T4 DNA ligase, T4 polynucleotide kinase, thermostable DNA polymerases for PCR, restriction endonucleases, exonuclease, and Gibson assembly kits were purchased from New England Biolabs. *E. coli* single-stranded DNA-binding protein (SSB) was supplied by USB. TS2126 RNA ligase I was generously provided by Dr. Kevin Ryan (City College of New York). Bio-Spin 6 desalting columns were obtained from BioRad. All T7 phage proteins and *E. coli* thioredoxin were prepared as described previously^29-33^; Uniprot IDs: T7 single-stranded DNA binding protein, gp2.5: (P03696 · SSB_BPT7), T7 DNA polymerase, gp5: (P00581 · DPOL_BPT7), T7 primase-helicase, gp4: (P03692 · HELIC_BPT7), *E. coli* Trx: (P0AA25 · THIO_ECOLI), *E. coli* SSB: (P0AGE0· SSB_ECOLI), TS2126 RNA ligase I: (C0HM52· RLIG_BPTS2).

#### DNA Polymerase Assay on Linear Templates

DNA polymerase assays were conducted using reaction mixtures containing a 90-mer DNA template annealed to a 5′ ^32^P-labeled common primer (5′-GAG GAG GGT CGG CGG-3′). The template consisted of either a random sequence (5′-ATC CTC CAC CCT CCC CAA CCT TTC CAT TCC TCA ACC CTT CAC CTC ACT TCA CTC CAC TAT ACC ACT CTC AAC CCT CCG CCG ACC CTC CTC-3′, with the underlined region annealed to the primer) or trinucleotide repeat sequences (5′-(CAG)_25_ CCG CCG ACC CTC CTC-3′ or 5′-(CTG)_25_ CCG CCG ACC CTC CTC-3′).

Reactions were performed in Buffer A (40 mM Tris-HCl, pH 7.5, 50 mM potassium glutamate, 10 mM MgClL, 5 mM DTT) with 500 μM of each dNTP, 100 nM primer/template, and the indicated amount of T7 DNA polymerase (a complex of T7 gene 5 protein with *E. coli* thioredoxin). Reactions were carried out at 37°C for 3 minutes. For reactions containing T7 gp2.5 or *E. coli* SSB, 5 μM of each protein was added to the mixture and incubated at room temperature for 5 minutes before the addition of DNA polymerase to ensure coating of the ssDNA region of the template. Reactions were terminated by adding sequencing gel loading buffer containing 98% formamide and 10 mM EDTA. Aliquots were loaded on a 20% denaturing polyacrylamide gel, and radioactive products were analyzed using phosphorimager scanning.

#### DNA Binding Assay

The binding of T7 gp2.5 or *E. coli* SSB to DNA was measured by incubating the indicated amounts of T7 gp2.5 or *E. coli* SSB with 3 nM 5′ ^32^P-labeled ssDNA in Buffer A for 10 minutes at room temperature. After the addition of a neutral dye, reaction products were loaded onto a 10% native gel and subjected to electrophoresis for 2 hours at 4°C. The dried gel was analyzed using phosphorimager scanning.

#### Preparation of Minicircle DNA

A minicircle containing an 86-mer random sequence (5′-gaccct ATC CTC CAC CCT CCC CAA CCT TTC CAT TCC TCA ACC CTT CAC CTC ACT TCA CTC CAC TAT ACC ACT CTC AAC CCT TCAcc) was constructed using a splint DNA and T4 DNA ligase as previously described^21,23,31^. Minicircles containing 25 trinucleotide repeats of either the CAG or CTG sequence (5′-gaccct (CAG)_25_ CCGcc and 5′-gaccct (CTG)_25_ CCGcc, respectively) were ligated using TS2126 RNA ligase I^22^. Minicircles were purified from the ligation reaction using an 8% denaturing polyacrylamide gel, and potential contamination with linear DNA was excluded by treatment with *E. coli* exonuclease I (New England Biolabs). All minicircles contained an 8-nucleotide common sequence (lowercase letters), including a T7 primase recognition site (ACCC). A 126-mer DNA of complementary sequence containing a 5′-T_40_ tail was annealed to the minicircle.

#### DNA Synthesis Assay on Minicircle Template

Leading strand DNA synthesis was measured by the incorporation of specific radioactive nucleotides: dGMP for the random sequence template, dAMP for the CTG repeat template, and dTMP for the CAG repeat template. Conversely, lagging strand synthesis was determined by measuring the incorporation of dCMP for the random sequence template, dTMP for the CTG repeat template, and dAMP for the CAG repeat template. Reactions were performed in Buffer A with 300 μM ATP and CTP, 500 μM of each dNTP, and the appropriate [α-^32^P] dNTP for leading or lagging strand synthesis. The reaction mixtures contained 10 nM minicircle template, 80 nM T7 DNA polymerase, 60 nM T7 gene 4 primase-helicase, and 4 μM T7 gene 2.5 protein. After incubation at 30°C, reactions were terminated at the indicated times by adding EDTA to a final concentration of 20 mM. Aliquots were spotted on DE-81 filter paper, washed three times with 300 mM ammonium formate (pH 8.0), and dried. The amount of ^32^P labeled deoxynucleoside monophosphate incorporated into DNA was determined by liquid scintillation counting.

Reaction products were also analyzed on a 0.8% alkaline agarose gel. After passing through a Bio-Spin 6 desalting column to remove unincorporated radioactive dNTPs, samples were heated at 90°C for 3 minutes before loading onto the gel. Electrophoresis was conducted at room temperature overnight, and the gel was dried and analyzed by autoradiography.

#### Nucleotide Hydrolysis Assay

Inorganic phosphate released from nucleotide hydrolysis was quantified using a malachite green assay^34^. Reactions contained 50 mM Tris-HCl (pH 7.4), 50 mM potassium glutamate, 500 µM dNTPs, 5 mM DTT, 100 nM gene 4 primase-helicase, 90 nM T7 DNA polymerase, and 10 nM minicircle DNA template. Reactions were quenched 1:1 with 50 mM EDTA, and 0.25 volumes of freshly prepared malachite green color solution (0.1% (w/v) malachite green oxalate (Thermo Scientific), 4.7 N H_2_SO_4_ (Sigma-Aldrich), 1.5% (w/v) ammonium molybdate tetrahydrate (Thermo Scientific), and 0.2% (v/v) Tween-20) was added. After 15 minutes at room temperature, absorbance was measured at 620 nm, and phosphate concentration was determined using KH_2_PO_4_ standards prepared in the same buffer.

#### Introduction of Trinucleotide Repeat Sequences into the T7 Genome

The 86-mer sequences (random or trinucleotide-repeat) used for minicircle construction were inserted into the T7 genome by replacing the non-essential gene 5.3 through homologous recombination. Plasmids for recombination contained the *trxA* gene for *E. coli* thioredoxin plus the test sequences, flanked by upstream and downstream regions of T7 gene 5.3. Three DNA fragments (a 440-bp region upstream of T7 gene 5.3, the thioredoxin gene followed by the 86-mer sequence, and a 520-bp region downstream of T7 gene 5.3) were combined using a two-step mega primer PCR with Q5 DNA polymerase. The DNA was inserted at a *SmaI* site of a pUC19 vector using a Gibson assembly kit. Notably, DNA containing 25 CTG repeats was prepared using a primer with the full 25 repeats, as shorter primers failed due to secondary structure formation during PCR. After infecting *E. coli* containing the plasmid with wild-type T7 phage, the resulting lysate was plated on a bacterial lawn of an *E. coli* strain lacking the thioredoxin gene (*TrxA*). Recombinant phages acquiring the thioredoxin gene were selected, and the integrity of the recombinant T7 genome was verified by analyzing the DNA sequence downstream of the thioredoxin gene. T7 DNA was amplified using diluted phage solution as a template in a PCR with Taq DNA polymerase.

## Data availability

All data presented are contained in the article and is available upon request from Alfredo J. Hernandez, Tufts University.

## Supporting information

This article does not contain supporting information.

## Acknowledgments

We thank members of the Hernandez laboratory and the Tufts Department of Biology for useful discussions.

## Funding and additional information

This work was supported by startup funds provided to A.J.H. by Tufts University.

## References

1. Khristich, A.N., and Mirkin, S.M. (2020). On the wrong DNA track: Molecular mechanisms of repeat-mediated genome instability. J Biol Chem 295, 4134–4170. 10.1074/jbc.REV119.007678.

2. Khristich, A.N., Armenia, J.F., Matera, R.M., Kolchinski, A.A., and Mirkin, S.M. (2020). Large-scale contractions of Friedreich’s ataxia GAA repeats in yeast occur during DNA replication due to their triplex-forming ability. Proc Natl Acad Sci U S A 117, 1628–1637. 10.1073/pnas.1913416117.

3. Budworth, H., and McMurray, C.T. (2013). Bidirectional transcription of trinucleotide repeats: roles for excision repair. DNA Repair (Amst) 12, 672–684. 10.1016/j.dnarep.2013.04.019.

4. Lin, Y.H., Chang, C.C., Wong, C.W., and Teng, S.C. (2009). Recruitment of Rad51 and Rad52 to short telomeres triggers a Mec1-mediated hypersensitivity to double-stranded DNA breaks in senescent budding yeast. PLoS One 4, e8224. 10.1371/journal.pone.0008224.

5. Cleary, J.D., Pattamatta, A., and Ranum, L.P.W. (2018). Repeat-associated non-ATG (RAN) translation. J Biol Chem 293, 16127–16141. 10.1074/jbc.R118.003237.

6. McMurray, C.T. (2010). Mechanisms of trinucleotide repeat instability during human development. Nat Rev Genet 11, 786–799. 10.1038/nrg2828.

7. Moore, H., Greenwell, P.W., Liu, C.P., Arnheim, N., and Petes, T.D. (1999). Triplet repeats form secondary structures that escape DNA repair in yeast. Proc Natl Acad Sci U S A 96, 1504–1509. 10.1073/pnas.96.4.1504.

8. Kunkel, T.A., Patel, S.S., and Johnson, K.A. (1994). Error-prone replication of repeated DNA sequences by T7 DNA polymerase in the absence of its processivity subunit. Proc Natl Acad Sci U S A 91, 6830–6834.

9. Thomas, D.C., Nguyen, D.C., Piegorsch, W.W., and Kunkel, T.A. (1993). Relative probability of mutagenic translesion synthesis on the leading and lagging strands during replication of UV-irradiated DNA in a human cell extract. Biochemistry 32, 11476–11482. 10.1021/bi00094a002.

10. Pearson, C.E., Tam, M., Wang, Y.H., Montgomery, S.E., Dar, A.C., Cleary, J.D., and Nichol, K. (2002). Slipped-strand DNAs formed by long (CAG)*(CTG) repeats: slipped-out repeats and slip-out junctions. Nucleic Acids Res 30, 4534–4547. 10.1093/nar/gkf572.

11. Fouche, N., Ozgur, S., Roy, D., and Griffith, J.D. (2006). Replication fork regression in repetitive DNAs. Nucleic Acids Res 34, 6044–6050. 10.1093/nar/gkl757.

12. Cleary, J.D., Nichol, K., Wang, Y.H., and Pearson, C.E. (2002). Evidence of cis-acting factors in replication-mediated trinucleotide repeat instability in primate cells. Nat Genet 31, 37–46. 10.1038/ng870.

13. Nelson, S.W., and Benkovic, S.J. (2010). Response of the bacteriophage T4 replisome to noncoding lesions and regression of a stalled replication fork. J Mol Biol 401, 743–756. 10.1016/j.jmb.2010.06.027.

14. Barry, J., Wong, M.L., and Alberts, B. (2019). In vitro reconstitution of DNA replication initiated by genetic recombination: a T4 bacteriophage model for a type of DNA synthesis important for all cells. Mol Biol Cell 30, 146–159. 10.1091/mbc.E18-06-0386.

15. Benkovic, S.J., and Spiering, M.M. (2017). Understanding DNA replication by the bacteriophage T4 replisome. J Biol Chem 292, 18434–18442. 10.1074/jbc.R117.811208.

16. Kulczyk, A.W., and Richardson, C.C. (2016). The Replication System of Bacteriophage T7. Enzymes 39, 89–136. 10.1016/bs.enz.2016.02.001.

17. Lee, S.J., and Richardson, C.C. (2011). Choreography of bacteriophage T7 DNA replication. Curr Opin Chem Biol 15, 580–586. 10.1016/j.cbpa.2011.07.024.

18. Patel, S.S., Wong, I., and Johnson, K.A. (1991). Pre-steady-state kinetic analysis of processive DNA replication including complete characterization of an exonuclease-deficient mutant. Biochemistry 30, 511–525.

19. Meyer, R.R., and Laine, P.S. (1990). The single-stranded DNA-binding protein of Escherichia coli. Microbiol Rev 54, 342–380. 10.1128/mr.54.4.342-380.1990.

20. Hernandez, A.J., and Richardson, C.C. (2019). Gp2.5, the multifunctional bacteriophage T7 single-stranded DNA binding protein. Semin Cell Dev Biol 86, 92–101. 10.1016/j.semcdb.2018.03.018.

21. Lee, J., Chastain, P.D., 2nd, Kusakabe, T., Griffith, J.D., and Richardson, C.C. (1998). Coordinated leading and lagging strand DNA synthesis on a minicircular template. Mol Cell 1, 1001–1010. S1097-2765(00)80100-8 [pii].

22. Blondal, T., Thorisdottir, A., Unnsteinsdottir, U., Hjorleifsdottir, S., Aevarsson, A., Ernstsson, S., Fridjonsson, O.H., Skirnisdottir, S., Wheat, J.O., Hermannsdottir, A.G., et al. (2005). Isolation and characterization of a thermostable RNA ligase 1 from a Thermus scotoductus bacteriophage TS2126 with good single-stranded DNA ligation properties. Nucleic Acids Res 33, 135–142. 10.1093/nar/gki149.

23. Lee, J., Chastain, P.D., 2nd, Griffith, J.D., and Richardson, C.C. (2002). Lagging strand synthesis in coordinated DNA synthesis by bacteriophage t7 replication proteins. J Mol Biol 316, 19–34. 10.1006/jmbi.2001.5325S0022283601953252 [pii].

24. Kusakabe, T., and Richardson, C.C. (1997). Template recognition and ribonucleotide specificity of the DNA primase of bacteriophage T7. J Biol Chem 272, 5943–5951.

25. Hartenstine, M.J., Goodman, M.F., and Petruska, J. (2000). Base stacking and even/odd behavior of hairpin loops in DNA triplet repeat slippage and expansion with DNA polymerase. J Biol Chem 275, 18382–18390. 10.1074/jbc.275.24.18382.

26. Stano, N.M., Jeong, Y.J., Donmez, I., Tummalapalli, P., Levin, M.K., and Patel, S.S. (2005). DNA synthesis provides the driving force to accelerate DNA unwinding by a helicase. Nature 435, 370–373. 10.1038/nature03615.

27. Wong, I., Patel, S.S., and Johnson, K.A. (1991). An induced-fit kinetic mechanism for DNA replication fidelity: direct measurement by single-turnover kinetics. Biochemistry 30, 526–537.

28. Brown, R.E., and Freudenreich, C.H. (2021). Structure-forming repeats and their impact on genome stability. Curr Opin Genet Dev 67, 41–51. 10.1016/j.gde.2020.10.006.

29. Tabor, S., and Richardson, C.C. (1985). A bacteriophage T7 RNA polymerase/promoter system for controlled exclusive expression of specific genes. Proc Natl Acad Sci U S A 82, 1074–1078.

30. Lee, S.J., and Richardson, C.C. (2001). Essential lysine residues in the RNA polymerase domain of the gene 4 primase-helicase of bacteriophage T7. J Biol Chem 276, 49419–49426. 10.1074/jbc.M108443200M108443200 [pii].

31. Hernandez, A.J., Lee, S.J., and Richardson, C.C. (2016). Primer release is the rate-limiting event in lagging-strand synthesis mediated by the T7 replisome. Proc Natl Acad Sci U S A 113, 5916–5921. 10.1073/pnas.1604894113.

32. Rezende, L.F., Hollis, T., Ellenberger, T., and Richardson, C.C. (2002). Essential amino acid residues in the single-stranded DNA-binding protein of bacteriophage T7. Identification of the dimer interface. J Biol Chem 277, 50643–50653. 10.1074/jbc.M207359200M207359200 [pii].

33. Lee, S.J., Ferguson, C., Urbano, S., Lee, J., Jeong, P., Cheela, M., Mitsunobu, H., Zhu, B., Prajapati, A., Richardson, C.C., and Hernandez, A.J. (2025). Mechanism of Annealing of Complementary DNA Strands by the Single-Stranded DNA Binding Protein of Bacteriophage T7. Biochemistry 64, 155s0–1559. 10.1021/acs.biochem.4c00730.

34. Baykov, A.A., Evtushenko, O.A., and Avaeva, S.M. (1988). A malachite green procedure for orthophosphate determination and its use in alkaline phosphatase-based enzyme immunoassay. Anal Biochem 171, 266–270. 10.1016/0003-2697(88)90484-8.

